# Galaxy Training: A Powerful Framework for Teaching!

**DOI:** 10.1101/2022.06.02.494505

**Authors:** Saskia Hiltemann, Helena Rasche, Simon Gladman, Hans-Rudolf Hotz, Delphine Larivière, Daniel Blankenberg, Pratik D. Jagtap, Thomas Wollmann, Anthony Bretaudeau, Nadia Goué, Timothy J. Griffin, Coline Royaux, Yvan Le Bras, Subina Mehta, Anna Syme, Frederik Coppens, Bert Droesbeke, Nicola Soranzo, Wendi Bacon, Fotis Psomopoulos, Cristóbal Gallardo-Alba, John Davis, Melanie Christine Föll, Matthias Fahrner, Maria A. Doyle, Beatriz Serrano-Solano, Anne Fouilloux, Peter van Heusden, Wolfgang Maier, Dave Clements, Florian Heyl, Björn Grüning, Bérénice Batut, the Galaxy Training Network

## Abstract

There is an ongoing explosion of scientific datasets being generated, brought on by recent technological advances in many areas of the natural sciences. As a result, the life sciences have become increasingly computational in nature, and bioinformatics has taken on a central role in research studies. However, basic computational skills, data analysis and stewardship are still rarely taught in life science educational programs [1], resulting in a skills gap in many of the researchers tasked with analysing these big datasets. In order to address this skills gap and empower researchers to perform their own data analyses, the Galaxy Training Network (GTN) has previously developed the Galaxy Training Platform (https://training.galaxyproject.org); an open access, community-driven framework for the collection of FAIR training materials for data analysis utilizing the user-friendly Galaxy framework as its primary data analysis platform [2].

Since its inception, this training platform has thrived, with the number of tutorials and contributors growing rapidly, and the range of topics extending beyond life sciences to include topics such as climatology, cheminformatics and machine learning. While initially aimed at supporting researchers directly, the GTN framework has proven to be an invaluable resource for educators as well. We have focused our efforts in recent years on adding increased support for this growing community of instructors. New features have been added to facilitate the use of the materials in a classroom setting, simplifying the contribution flow for new materials, and have added a set of train-the-trainer lessons. Here, we present the latest developments in the GTN project, aimed at facilitating the use of the Galaxy Training materials by educators, and its usage in different learning environments.

## Introduction

Education is a fundamental human right (e.g. the Universal Declaration of Human Rights (UDHR), the International Covenant on Economic, Social and Cultural Rights (ICESCR)). Indicators of the achievement of education as a right are outlined in the ICESCR, and further developed into what is known as the “4 As framework” [3], which specifies Availability, Accessibility, Acceptability and Adaptability as essential metrics. The 4 As, therefore, provide a concrete set of ideals to strive towards in any global educational endeavor. The goals of the Galaxy Training Network (GTN; https://training.galaxyproject.org) and the Galaxy Project are well aligned with the 4 As.

Galaxy [4] is an open source platform for accessible, reproducible, and transparent computational research, driven by an inclusive and diverse worldwide community. Researchers are able to access a wealth of tools (*>*8,500 tools in the Galaxy ToolShed [5] as of March 2022), datasets, and high-performance compute resources, through a standard web browser, without requiring informatics expertise. However, comprehensive training is still required to adequately understand the data analyses and to accurately interpret the results. A survey of 704 NSF-BIO-funded investigators revealed that training topics were the top 3 of 13 most unmet data analysis needs, above HPC/cloud facilities, workflows/pipelines analysis or storage facilities needs [6].

To address this large demand for training, the GTN was founded to provide learners with free online training materials, connect them with local trainers, and help promote open data analysis practices worldwide [2]. These training materials are openly developed, maintained and community-reviewed via GitHub (https://github.com/galaxyproject/training-material). The materials cover an increasing spread of topical domains, such as life sciences, computational chemistry, climate sciences, data visualisation, statistics, and machine learning.

The GTN website has seen a steady increase in usage since its creation (Figure 1D), with in average 28,000+ visits per month. Since September 2018, 1,800+ individual tutorial feedback forms have been submitted with a high satisfaction rate (more than 88% had a satisfaction rate of 4 or 5, per Figure 2D).

**Fig 1.**
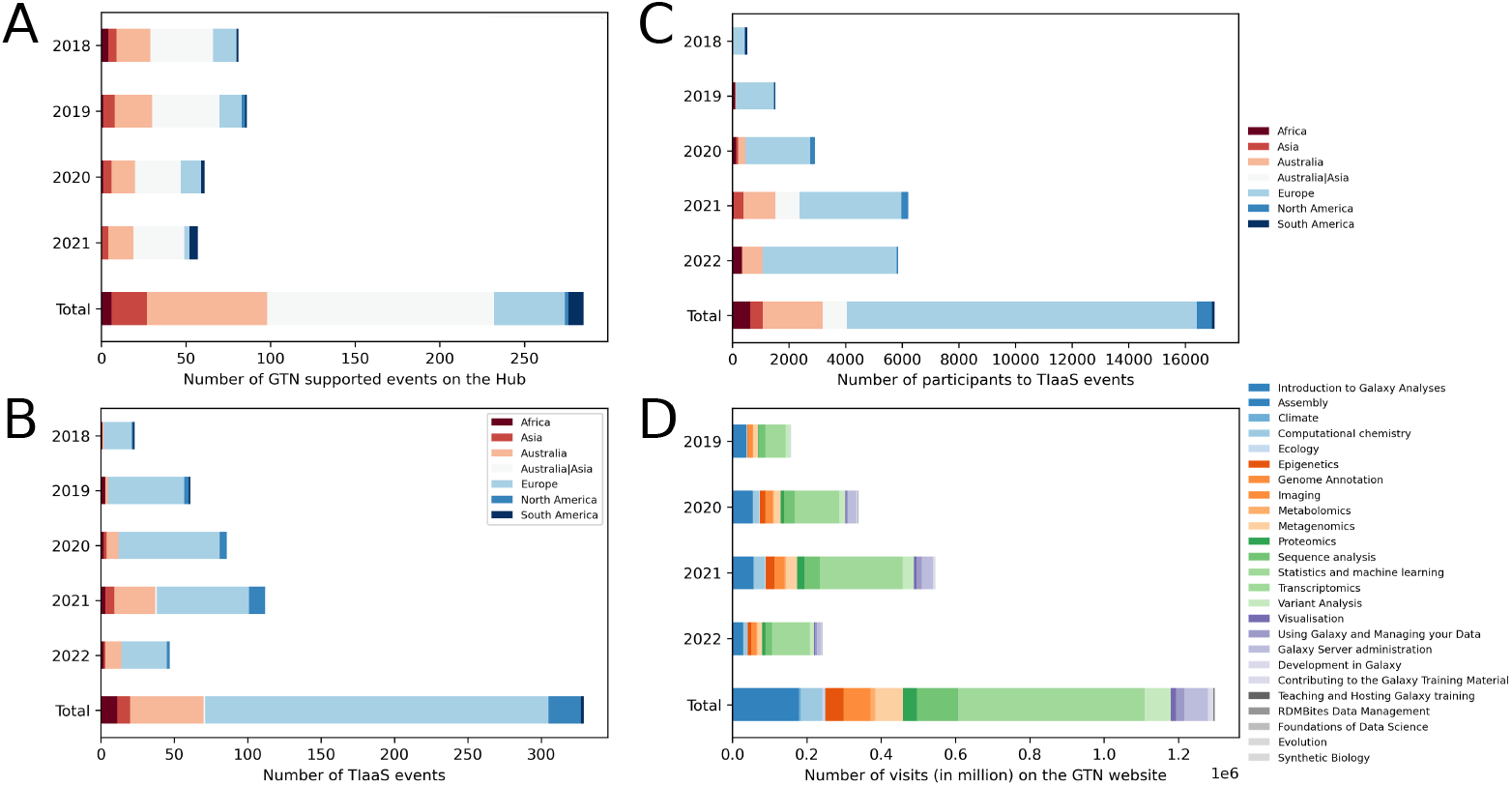
Statistics about training events organized by the Galaxy community and visits of the GTN website over years. A: Number of events supported GTN materials, registered on the Galaxy Community Hub website [9], per year between 2018 and 2021. B: Number of Training Infrastructure as a Service (TIaaS) events per year on Galaxy Europe (collected in May 2022). C: Number of participants to TIaaS events per year on UseGalaxy.* (collected in May 2022). D: Number of visits per year on the GTN website and per topics, initially tracked by Google Analytics and later with Plausible (collected in May 2022). The latest usage statistics are publicly available from https://plausible.galaxyproject.eu/training.galaxyproject.org

**Fig 2.**
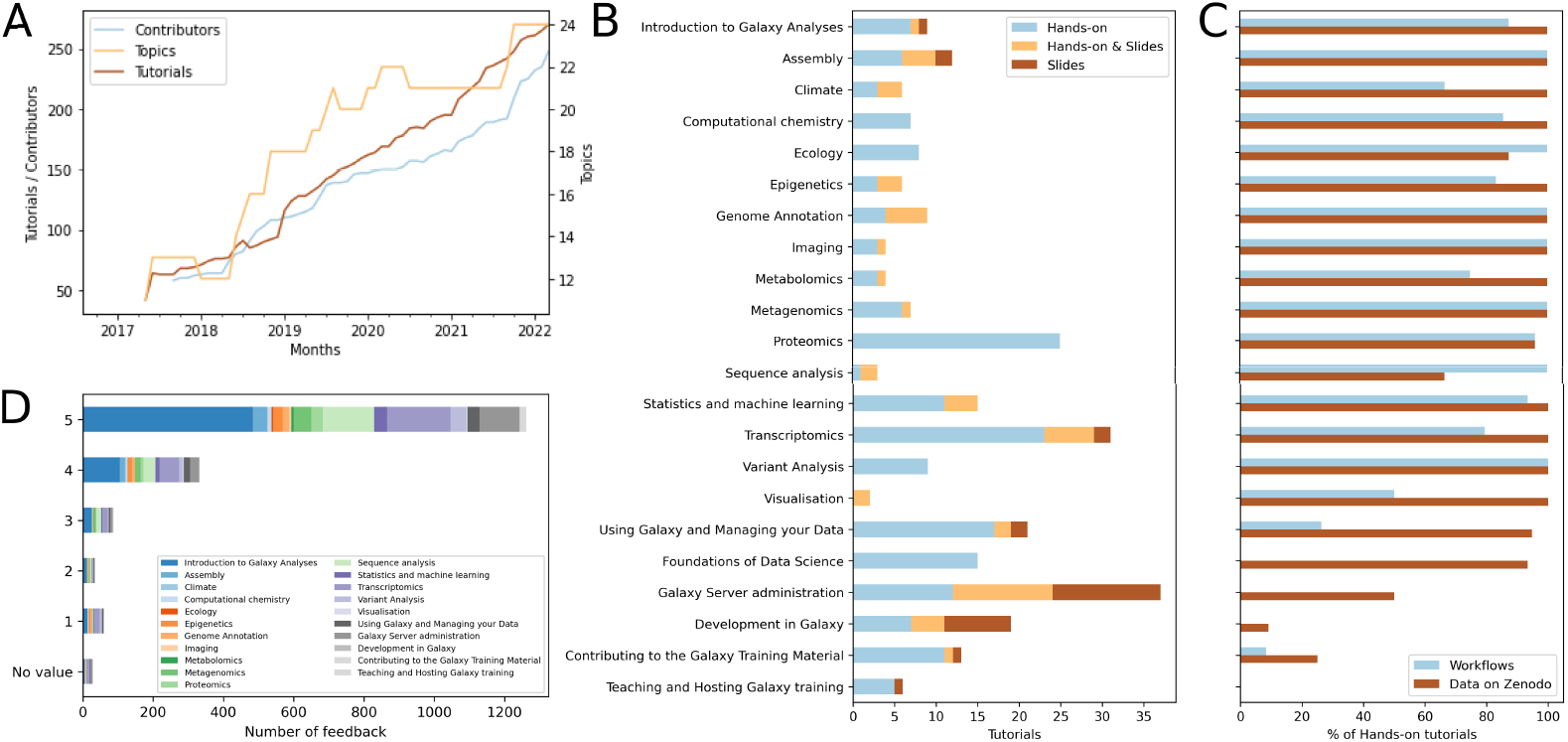
Content of material available on the GTN and feedback from learners. A: Evolution of number of topics, tutorials and contributors over the months between 2017 and 2022. B: Number and type of tutorials per topics available on the GTN on April 2022. The latest statistics are publicly available from https://training.galaxyproject.org/stats. C: Type of supporting materials for tutorials per topics available on the GTN on April 2022. D: Score of the embedded feedback in the tutorials per topics. Three questions are asked in the form: “How much did you like this tutorial?” (from 1 (bad) to 5 (great)), “What did you like?”, “What could be improved?”. The latest feedback results are publicly available from https://training.galaxyproject.org/feedback

By utilizing Galaxy as its primary data analysis platform, the GTN also addresses the clear demand for easily accessible compute resources for research and training in life sciences [6]. Galaxy is an ideal platform for teaching, since it takes care of all the heavy lifting behind the scenes, such as installing software and managing the compute infrastructure, leaving learners free to focus on the scientific analysis concepts at hand, rather than on the details of the technical implementations.

By using one of the worldwide public Galaxy servers, learners and educators alike gain free access to high-performance compute resources without any of the systems administration burden. These include the UseGalaxy.* servers with Galaxy Main (https://usegalaxy.org), Galaxy Europe (https://usegalaxy.eu), and Galaxy Australia (https://usegalaxy.org.au), but also 160+ others [7]. Therefore, over the past years, Galaxy has become increasingly attractive as a training platform for instructors around the world (Figure 1A). For example, between June 2018 and May 2022, UseGalaxy.* servers have been used for 330+ registered training events reaching 17,000+ learners. In addition to these short-running training events, GTN tutorials have also been included as part of formal undergraduate and graduate courses (see User Stories section for more details).

Given this observation of the value of Galaxy and the GTN in teaching activities, we have aimed to optimize the GTN framework to support educators in the development and reuse of training materials, e.g. by integrating the FAIR (Findable, Accessible, Interoperable, Reusable) principles for training materials [8] directly into the framework. The Galaxy Training Network and its framework have then grown significantly over the past years (Figure 1). In this paper, we describe the aims and objectives of the GTN and its framework, and highlight some recent efforts by the GTN community to expand and improve the project. We also present how this project supports both educators and lesson developers to bring availability, accessibility, acceptability, and adaptability to bioinformatics education for trainers and learners alike. Finally, we showcase some example user stories of how the GTN materials are being used across all levels of education.

## Results

### GTN Overview: Materials and Features

The GTN Materials are continually growing, both in terms of the number of tutorials as well as its community (Figure 2A). The GTN Materials in May 2022 has 260+ tutorials covering 23 topics (17 scientific, 6 technical), developed by over 260+ contributors (Figure 2A).

#### Tutorials

Tutorials are typically constructed around a real-world *research story*, e.g. a published journal article describing an analysis workflow or dataset. The tutorial starts by introducing the relevant scientific background, and then proceeds to describe the analysis, providing details on the relevant scientific and computational concepts involved at each step. The tutorials are composed of alternating theory and practical sections, interspersed with formative assessment questions and exercises, and may be supported by a set of introductory slides (Figure 3). Our tutorials are self-contained; everything needed to complete them is bundled with the tutorial; this includes all the datasets, lecture slides, videos, the workflows, and the list of public Galaxy servers supporting the tutorial. No software installation is needed to follow these tutorials; the only technical requirement is access to internet.

**Fig 3.**
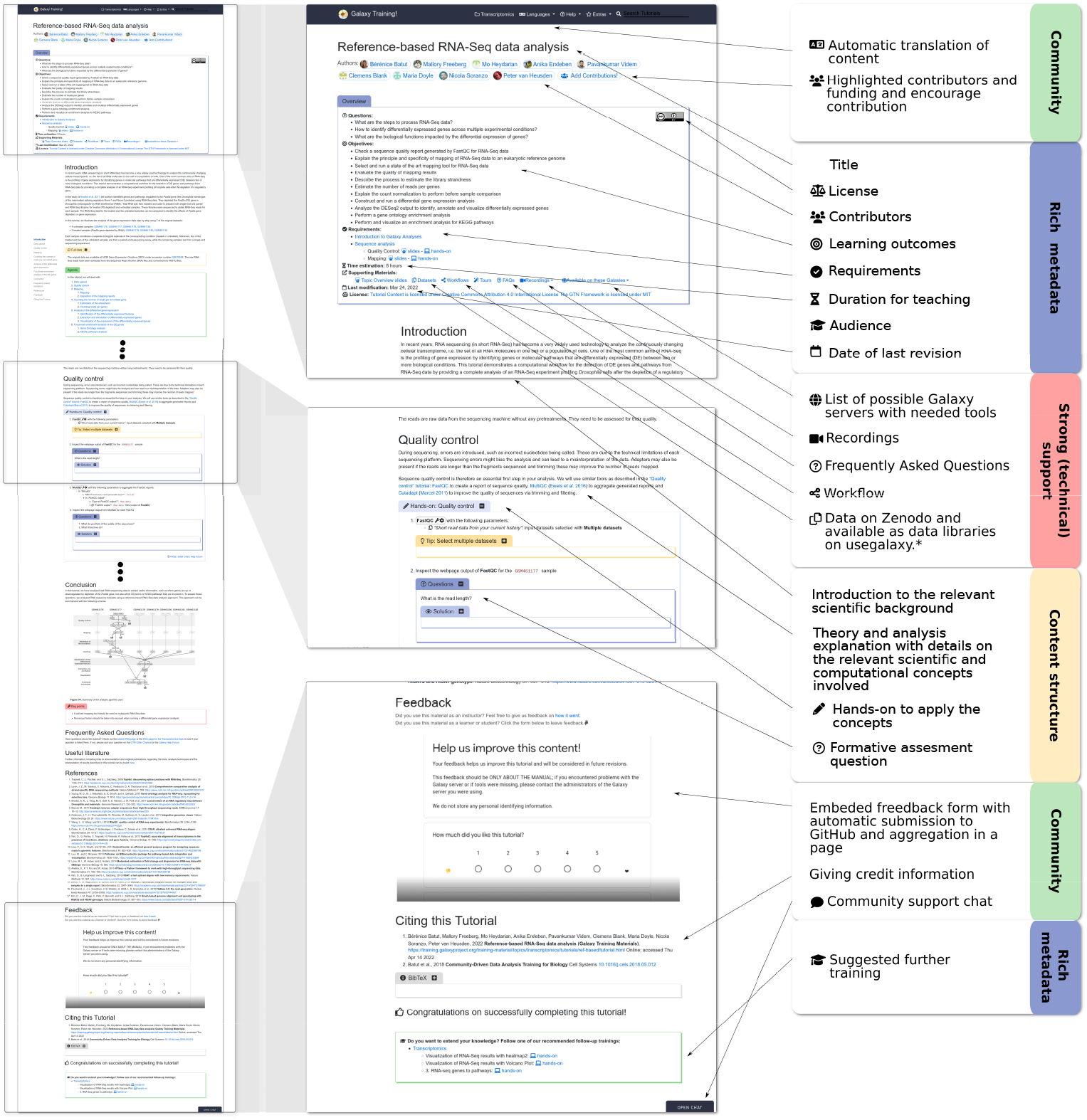
Structure of a GTN tutorial. Title and authors are listed, followed by an overview box containing metadata about the tutorial (target audience, learning objectives, prerequisites, supporting materials, time estimate, etc). The tutorial content itself is a mix of theory (text) and so-called “hands-on” boxes describing practical steps to be performed in Galaxy. Question and answer boxes may be added at any point in the tutorial to allow students to self-evaluate as they progress. The end of a tutorial provides a box with take-home messages, references and suggestions for further reading and follow-up tutorials. Additionally, a feedback form is embedded at the end of every tutorial. Access to FAQs, support channels, page view statistics, and translated versions of the materials are provided via the top menu of the webpage.

A survey of trainers identified a desire of increased interactivity between tutorials and Galaxy. We have developed a new Galaxy feature “Tutorial Mode” which allows following the training materials directly in Galaxy. With that mode activated, hands on steps are now interactive and clicking tools brings you directly to the tool in Galaxy.

#### Recording and automated video lectures

In order to further expand the formats in which our training materials are available, several tutorials have been recorded by community members, are available on GTN Training Video Library [10] and accessible directly from the tutorial itself. Recording a video is a laborious and time intensive activity. To save a significant amount of instructor time, we automatically generate videos based on lecture slide decks. The slides are narrated using automated text-to-speech (TTS), and the script is based on the speaker’s slide notes, which makes for an extremely easy video update process–updating the slides automatically re-records the videos.

#### Coding Tutorials

Jupyter and RStudio [11] can both be run within Galaxy in the form of an *interactive tool*, allowing for interactive tutorials that offer a combination of Galaxy-based analysis and coding-based R or Python analysis steps. In addition to covering a wide range of analyses in Galaxy, the GTN also supports coding-oriented tutorials in the form of Jupyter [12] and RMarkdown [13] notebooks which are automatically generated from the tutorial content.

#### FAIR Training

The GTN infrastructure has been developed in accordance with the FAIR principles for training materials [8] (Table S2) and following the 10 simple rules for collaborative lesson development as defined by Deveny et al. [14] (Table S1). Following these principles enables trainers and trainees to find, reuse, adapt, and improve the available tutorials.

### GTN for Educators

The GTN training platform as described before helps minimize the amount of time and effort required for instructors to prepare for and run their training courses and workshops, by providing tutorials and a complete training infrastructure.

#### Train-the-trainer (TtT) Tutorials

To support teachers and trainers, we have developed a dedicated topic in the GTN for TtT tutorials, covering all aspects of using the GTN materials in education. This includes best practices providing pedagogical, technical and logistical recommendations, accessible on the website with a series of tutorials (available in “Teaching and Hosting Galaxy training” category). For first-time instructors, the GTN has come together to collect different instructors’ experiences (“Training Philosophies” [15]).

#### Preparing a Workshop

During preparation of a workshop, organizers and instructors work together to identify relevant tutorials for their event. Our tutorials are roughly divided into “topics” such as *Transcriptomics* or *Climate Science*, with a search function available on the GTN website (https://training.galaxyproject.org). Within topics, multiple tutorials can be found. Each tutorial starts with a list of metadata such as learning objectives following Bloom’s taxonomy [16], prerequisites, time estimate or questions addressed by the tutorial (Figure 3). This but also tags of tutorials enables instructors to identify the best tutorials for their audience. Reusable slide decks with speaker notes are also available to introduce the topics prior to a tutorial.

To ensure the success of practical sessions, the tutorials rely on specific datasets (included in the tutorial metadata), and tools which the GTN analyzes in order to provide a list of compatible public Galaxy servers. Automated workflow testing [17] provides reassurances to instructors, letting them know that a given tutorial keeps working on their selected server.

We have added support for FAQs to be added to tutorials, in order to help instructors prepare their lesson. These FAQs typically cover common questions or frequently observed trainee mistakes encountered during a particular tutorial.

#### Preparing Workshop Infrastructure

A major challenge in computational workshop organization is the identification of affordable and reliable compute infrastructure. The needs of training infrastructure are significantly different from regular research; analysis steps must complete in relatively short time periods in order to not disrupt the flow of the lesson. The GTN uses small (sub-sampled) training datasets for tutorials in order to reduce the run time of the analysis step, but a second factor to consider is the waiting time of the compute infrastructure. In order to support training events, we developed Training Infrastructure as a Service (TIaaS) [18]. This free service provides a dedicated job queue for the participants of training events, in order to reduce waiting times on the cluster and ensure courses run smoothly and efficiently. Furthermore, the TIaaS service also provides instructors with a dashboard, enabling them to monitor the progress of the participants. To date, TIaaS has been widely used: 17, 000+ students were taught over the course of 330+ events between June 2018 and May 2022, with 65% using the GTN materials (Figure 1B,C).

#### Teaching a Workshop

During the workshop, instructors can introduce the topic using the available slide decks, supported by detailed speaker notes. After this theoretical introduction, instructors may either use a *live-demo* approach, guiding students through the tutorials in a step-by-step fashion, or alternatively let students work through the tutorials at their own pace while providing support. The tutorials are a fully self-contained teaching resource as described in Figure 3, and therefore, support both these training modalities.

#### Feedback from Instructors

In January 2020, we conducted a survey asking how trainers used the available training material and infrastructure. We received answers from 33 trainers, 88% of which had conducted a training event in the last 3 years. This sample consisted of a continuum of trainers, from occasional to seasoned trainers. Of those recently giving training events, 79% of them used GTN resources, and some even developed new GTN materials for the occasion.

The GTN tutorials are well liked and shared; 91% of respondents indicating they would recommend them. The use of GTN material allows them to have more time to focus on the fundamental principles of analyses and the specific needs of the trainees. Of the surveyed trainers, 69% are contributors to the GTN, and among those who are not, 55% declare that they plan to become one in the future.

### Contributing to the GTN

The GTN framework also aims to ease the burden of tutorial development, contribution and maintenance by offering a comprehensive set of tools and standards for tutorial authors.

#### Tutorials about Contributing

To get acquainted with the tutorial development process, a contributor can start by consulting a set of dedicated tutorials in the *Contributing* topic in the GTN. These tutorials provide a step-by-step guide to creating a tutorial in the GTN framework, covering everything from technical guides about the framework to pedagogical best practices and submission of tutorials to GitHub.

Figure 4 depicts the typical process a tutorial contributor will go through when developing a new tutorial. The first step is usually to develop the analysis workflow in Galaxy. The challenge is to identify or create suitable input datasets. The selected data must be informative enough to illustrate the meaning of a given analysis, but not too large as to require long waiting times for upload or processing during a training event. The selected data could be a toy dataset created from scratch, or (preferably) an informative subset of a real-life dataset, for example using a single chromosome of a whole-genome dataset.

**Fig 4.**
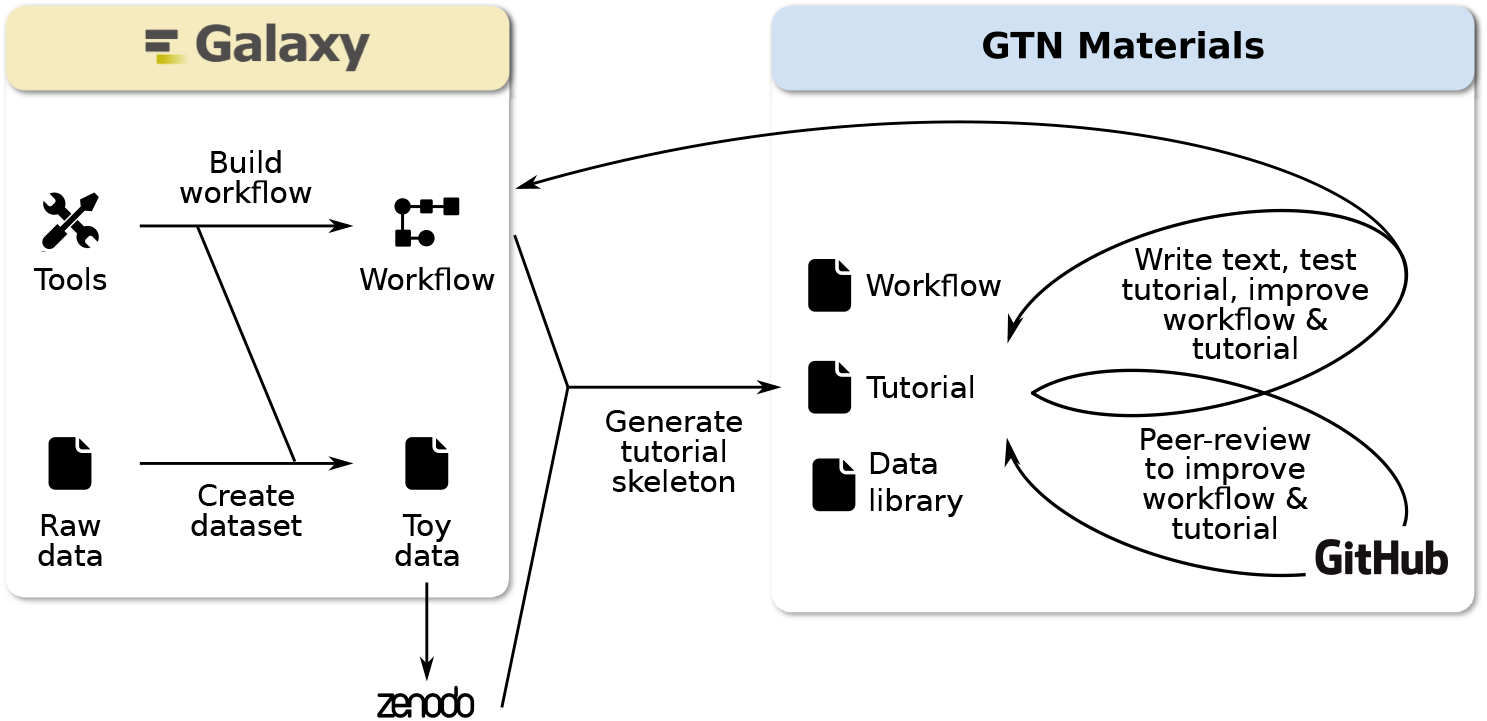
Typical process to create a new tutorial. Authors usually start by identifying suitable input datasets, and developing the workflow in Galaxy. Thus, a workflow can be automatically converted into a tutorial skeleton using the PTDK described in the next section. The tutorial is then tested, reviewed and finally merged into the GTN.

#### Planemo Tutorial Development Kit (PTDK)

To simplify the tutorial development process, we have created a *Tutorial Development Kit* within Planemo [19]. This automatically generates a tutorial skeleton from a Galaxy workflow as starting point for tutorial authors. This skeleton contains the overall structure of the tutorial: metadata section, example question boxes for formative assessments, instructions on how to proceed with adding scientific and pedagogical content and more important auto-generated hands-on boxes for every analysis tool in the workflow with the parameters to select-This process greatly reduces the development time, and allows tutorial creators to focus on the scientific content of the tutorial, rather than the technical details and style guidelines.

In order to further lower the contribution barrier, this *Planemo Tutorial Development Kit* has been encapsulated into a web service (https://ptdk.herokuapp.com/), where tutorial authors can provide a link to a public workflow on one of the UseGalaxy.* servers, and obtain the tutorial skeleton based on this workflow in Markdown, but also the workflow and data library file if a Zenodo link is provided. This removes the need for contributors to install and run Planemo locally. We have found that this approach saves time for not only the contributors but also for reviewers, since the tutorials contributed using this method adhere well to the GTN style guide.

#### Content reuse

To avoid duplication of content between tutorials, we have developed a set of modular tutorial components, called *snippets*, which can be easily reused across tutorials. For example, common tasks in Galaxy such as starting a workflow or creating a new history will be a part of most tutorials. Contributors can include these snippets at any place in a tutorial with a simple import statement. If instructions for this common task change, e.g. due to changes in the Galaxy user interface, the changes need to only be made in the snippet itself and will automatically propagate to any tutorial using them.

#### Tutorial Preview and Testing

GTN tutorials are written in Markdown format for ease of contribution, and subsequently converted to HTML web pages automatically by our framework. The HTML web page can be locally previewed if desired by using simple commands. Additionally, contributors who do not wish to install and run the GTN locally, can also generate a preview of their in-development tutorials online using GitPod [20]). We have integrated the GTN GitHub repository with GitPod, enabling contributors to obtain an online tutorial development environment, complete with online preview, with a single click of a button. GitPod has proven particularly useful for collaboration between multiple lesson developers with varying degrees of familiarity with git or GitHub.

In addition to a visual inspection of the generated web pages, our framework offers a suite of testing tools, allowing contributors to check that their tutorial meets the technical and style guidelines. These tests include checks of whether all required metadata is present, whether links within the tutorial are valid, and whether files are correctly formatted. This helps contributors and reviewers to quickly identify and correct potential problems with a tutorial.

#### Peer Review

Once a contributor is happy with their tutorial, they can create a *pull request* (PR) to the GTN GitHub repository (https://github.com/galaxyproject/training-material), where each contribution will then undergo automatic quality assurance tests and a peer-review process.

This review process is completely open, and any volunteers from the community may participate. For each topic within the GTN, we have encouraged several prominent community members who are regular contributors to act as topic *maintainers* that help safeguard the quality of the content in the topic, and are empowered to review, approve, and merge contributions. Maintainers and other reviewers will check the proposed contribution, both in terms of formatting and scientific content. They can make suggestions for updates, and start a discussion with the contributor(s). Typically, there will be two or three reviewers for any given PR. Based on these reviews, the contributor(s) will update the tutorial, and this cycle of update and review will continue until both contributor and reviewer are happy with the result. In case of disagreements, topic maintainers will decide on the path forward.

This open strategy for content creation and updates is paying off. Since the beginning of the project in the middle of 2016, over 2,500 PRs have been created to add or update tutorials.

The review process is quite fast: it takes 20 days between the opening of a pull request and its merge. Thanks to a fast turnaround, tutorials are regularly updated and iteratively improved over time, keeping the material relevant and of a high quality. Indeed, each tutorial undergoes (on average) a PR every quarter to update it. Some highly used tutorials, like the “Reference-based RNA-seq” tutorial, have received 100+ PRs over the last 3 years, on average one every 3 weeks.

#### Maintenance

Tutorials are dynamic entities; underlying analysis tools receive updates, new tools are developed, and the Galaxy interface itself changes regularly. As a result, tutorials must undergo regular updates as well to reflect the state-of-the-art in the scientific domain, and correctly reflect the latest available version of the tools and the Galaxy interface. Instructors preparing for their next workshop check the tutorials as preparation, and in the process, identify the places where the tutorial should be updated. If the instructors feel comfortable making these changes themselves, they can open a PR proposing the changes. If not, they can create an *issue* on GitHub to request the changes.

#### Feedback

Learner feedback is one of the most valuable resources to inform training improvements. An anonymous feedback form is embedded at the end of every tutorial, where users can provide feedback and make suggestions for enhancement. All feedback is transparently collected on the GTN website, enabling the community to view and address the feedback (Figure 2D).

### User Stories: GTN across all levels of education

#### GTN in Higher Education

Galaxy and the GTN are an integral part of Bioinformatics programs at the undergraduate level (e.g. Clermont Auvergne University, Texas A&M University, Avans Hogeschool, University of Freiburg, University of Frankfurt).

In these undergraduate courses, Galaxy is often used to teach the concept of data analysis, pipelines / workflows, and also reproducibility. Students learn the importance of the tool versions, as tools can change subtly or significantly over time and this may impact their analyses. Teachers can likewise showcase the evolution of algorithms over time using different versions of tools (e.g. bowtie and bowtie2) to help students understand the advances in the field. Without the need to spend resources on teaching command line skills and tool installation, the time can be fully used to introduce each step (data cleaning, data analysis and visualization) and to go into details of each tools and algorithms. Galaxy is also convenient to explain how data is structured (data types, metadata), and its connection to tools. Finally, Galaxy can be used to illustrate complete analytical workflows paying attention to input and output data.

#### GTN for Research Scientists

Postgraduate courses (e.g. Agrocampus Ouest, Rennes University, Brest University, Clermont Auvergne University, Station Biologique de Roscoff, Melbourne University, University of Freiburg) as well as internal bioinformatics short-format training sessions (e.g. Friedrich Miescher Institute, French Bioinformatics Institute, Erasmus MC, University of Freiburg) are relying on Galaxy as the main teaching tool. These courses are often directed at learners who are not yet comfortable working with the command line. However, we often notice that after getting familiar with each analysis step and understanding the meaning of parameters of each tool, some may feel confident to learn command line or scripting (possibly within Galaxy using RStudio or Jupyter notebook) and to move to command line environments.

GTN materials are also often used to provide supplemental training to later-career research scientists, for example at the Erasmus Medical Center (EMC), Earlham Institute, and as part of the Gallantries project [21]. These trainings often consist of short-running workshops, usually aimed at familiarizing researchers with novel analysis techniques. Since many of the GTN tutorials are centered around an analysis pipeline described in a recently published article, they are ideally suited to provide researchers with an update on the latest state-of-the art analysis pipelines in their domain. GTN tutorials are also frequently created as part of scientific publications by authors presenting a novel analysis method, as an additional form of documentation for the readers [22–27].

#### Training in underrepresented communities

Reliable internet access is often taken for granted by researchers. Bottlenecks in infrastructure may, however, cause significant issues for bioinformatics training events, as many students try to upload or download data at once. This is especially problematic in low or middle income countries (LMICs) where internet access may be intermittent, restricted or completely unavailable.

Trainers are often brought in from afar, meaning that the teaching takes places in –for the trainer– an unfamiliar setting, and on a strict time schedule limited by the return trip(s) of those involved. It is therefore important to avoid or minimise unforeseen delays caused by incompatibilities with local infrastructure, connectivity failures or unforeseen updates forcing new software to be downloaded or queries to remote servers to fail.

Based on these needs, the eBioKit [28] was developed: it is an assembly of open source or free-for-academic-use software, along with key databases and selected material for bioinformatics. It can be installed beforehand, brought on a portable server to the local training, and made available on the local network. Students can then access it through the network and work directly on the server, which avoids installation issues for the students, unforeseen updates of web services, or other failures. Galaxy has been a part of the eBioKit for most of its existence, specially in the eB3Kit which includes a Galaxy-based bioinformatics platform with a specialised workflow interface [29].

Various versions of the eBioKit has been used for over a decade by organisations such as EMBnet, H3Abionet, SANBio, and BECA/ILRI to train hundreds of researchers and bioinformatics trainers in LMICs.

Teaching based on Galaxy is also valuable in LMICs with limited internet access. The training datasets are all available online, either on the Galaxy server, in shared data libraries or in a third-party service such as Zenodo. Thus, the learners own (poor) internet connections are bypassed for these bandwidth-heavy tasks. The internet is only used to launch the analysis steps and for the teacher to monitor the progress of the analysis in Galaxy via TIaaS.

For cases where internet access is frequently absent, the GTN offers Docker [30] images, allowing for completely offline training. Every topic has its own Docker image, preconfigured with all the necessary tools, workflows, tours and data-libraries to complete the tutorials within that topic. A drawback of this solution is that all required compute resources to complete the tutorials must be available locally.

#### Citizen science and education

Thanks to its graphical user interface, Galaxy can also be used to introduce bioinformatics to a general audience and include them into scientific projects.

The Street Science Community [31] offers workshops to introduce biology, genomic sciences and bioinformatics to the public. For example, participants extract yeast DNA out of beer bottles and sequence it using a MinION, the Oxford Nanopore sequencing device. The generated sequencing data are then processed by participants inside Galaxy: uploading data, running a metagenomic analysis workflow, and visualizing results. Combining lab work and bioinformatics data analysis with Galaxy vividly demonstrates the challenges and possibilities that genomics brings to our society.

The GTN has also provided an opportunity for the ecology community to experiment with several ways to extend citizen science schemes, for instance, through two monitoring schemes from Vigie-Nature [32], French citizen science programs about common birds (Suivi Temporel des Oiseaux Communs) and bats (Vigie-Chiro). Volunteers use Galaxy to analyze data they gathered, as researchers would do, through user-friendly tools and workflows, documented in GTN. Such solutions increase the motivation of participants, especially volunteers, as they can analyze and visualize data about species and ecological systems they monitor, sometimes for several years.

#### Training in the COVID-19 era: Remote Learning

Due to the global COVID-19 pandemic, many educators have found themselves forced to change the modality of their training activities to completely virtual events. Galaxy and the GTN cater to remote learners and teachers with a set of features to facilitate the online learning process. It provides easy access to data and the possibility to share the progress and achievements, both student-to-student and student-to-instructor [33]. This has been extensively tested during the COVID-19 pandemic. For example, the GTN community organized the GTN Smörgåsbord events [34], global, 5 day, 24/7 events. These were fully asynchronous events, where instructors pre-recorded GTN tutorials, which the 2,000+ registered participants could work through at their own pace, with 120+ instructors available online for support on Slack across all time zones. This approach has been repeated for the Galaxy Community Conference training week [35], as well as various other virtual training events, including (but not limited to):

1. a 5-day workshop on “Machine Learning using Galaxy” organized by the Galaxy Europe team in June 2020, with 400+ registrations and 200+ participants on the first day [36]
2. A 5-day plant transcriptomics workshop organized by the Galaxy Europe team [37]
3. A Microbiome Informatics 5-day workshop organized by the Galaxy-P team with Galaxy India
4. A SARS-CoV-2 data analysis workshop [38]
5. A case study in COVID19 data analysis at the Great Lakes Bioinformatics Conference (GLBIO 2021)
6. Metatranscriptomics analysis using microbiome RNA-seq data in Galaxy
7. The Spanscriptomics workshop, a pilot study into using Spanish-translated GTN tutorials to teach single-cell analysis to native Spanish speakers.

#### Hybrid learning in geographically sparse locations

Even before the pandemic, learners in remote areas have faced significant barriers to travelling to in-person training events. In such areas, a so-called *hybrid training* [39] approach offers a solution. In such an approach, learners gather in classrooms in several geographically distinct locations. The training is live-broadcast to each of these locations, and communication with the instructor happens in real-time.

GTN resources have been successfully used for such hybrid training events, for example by Galaxy Australia, which organized a large number of hybrid training events, with up to 11 satellite classrooms across the region participating, and reaching over 800 learners [39]. The Gallantries project [21] has successfully applied a similar approach in the European region.

## Conclusion

From the start of the Galaxy project, education has been an important focus. Since 2018, more than 350 training events have been carried out. Recipients of these trainings included undergraduate and postgraduate students, research scientists, underrepresented communities and citizens. The foundation for most of these events were the Galaxy tutorials provided by the GTN using its well-structured and easy to maintain Galaxy-based training resource.

Since then the GTN has grown steadily and today more than 260 tutorials developed by more than 260 contributors across 23 topics are available. The GTN infrastructure is in accordance with the 4 As framework and the FAIR principles. It furthermore enables remote learning also in settings with intermittent, restricted or unavailable internet access. Teachers are empowered to prepare and run their trainings by specialized training resources for educators and technical features such as training infrastructure as a service (TIaaS). Also, contribution to GTN has been facilitated by the creation of dedicated tutorials and technical tools such as the Planemo tutorial development kit (PTDK) and GitPod.

To support the instructors and build a community of instructors, we are also collaborating with the Gallantries project and ELIXIR to build a Train the Trainer (TtT) program and a mentoring program for instructors. For contributors, we are also working with publishers to implement a system for publication of tutorial via article with limited extra work. Following the feedback from learners, instructors and contributors, new features are currently in development.

The Galaxy Training Network is a collaborative effort to develop and maintain state-of-the-art Galaxy based training in the life-sciences and beyond.

## Acknowledgments

This project would not be possible without the tireless efforts and valuable contributions from the world-wide GTN community [40].

This project has received funding from the Erasmus+ programme of the European Union (2020-1-NL01-KA203-064717). SH, HR acknowledge support from the European Union’s Horizon 2020 research and innovation programme under grant agreement No 825775, DB from NIH NHGRI 5U24HG006620 and NIH NCI 5U24CA231877, PDJ from ASM-IUSSTF Visiting Teaching Fellowship, NS from the Biotechnology and Biological Sciences Research Council (BBSRC), part of UK Research and Innovation, Core Capability Grant BB/CCG1720/1 and the National Capability BBS/E/T/000PR9814, ACF from the European Union’s Horizon 2020 programme under grant agreement No 857652 (EOSC-Nordic) and No 101017501 (RELIANCE). PVH from South African Research Chairs Initiatives of the Department of Science and Technology and National Research Foundation of South Africa grant UID 64751 and South African Medical Research Council flagship program MRC-RFA-UFSP-01-2013/COMBAT-TB, BB, BG, WM from the German Federal Ministry of Education and Research BMBF grant 031 A538A de.NBI-RBC.

## Availability

All training materials mentioned in this paper are stored on GitHub (https://github.com/galaxyproject/training-material/), freely available online (https://training.galaxyproject.org/), and licensed under a Creative Commons Attribution 4.0 International License.

The infrastructure behind the project is stored on GitHub (https://training.galaxyproject.org/) and licensed under a MIT License.

Most figures in this paper have been generated using Jupyter notebook available on the GitHub repository for this paper (https://github.com/galaxyproject/GTN-community-paper-2020).

## Supplementary Materials

**S1 Table.** Ten Simple Rules for Collaborative lesson development. Implementation of the “Ten simple rules for collaborative lesson development” [14] in the GTN.

| Rules | Implementation in the GTN framework |
| --- | --- |
| Clarify audience | Tutorial metadata includes level indicators (introductory, intermediate, advanced) and a list of prerequisite tutorials as recommended prior knowledge. This information is rendered at the top of each tutorial. |
| Make lessons modular | Development of small tutorials linked together via learning paths |
| Teach best practice lesson development | A topic *Contributing to the Galaxy Training Material* including 10 tutorials describing how to create new content. Furthermore, quarterly online collaboration fest (CoFests) are organized, where contributors can get direct support. Development of a Train the Trainer program and a mentoring program for instructors, in which lesson development is taught |
| Encourage and empower contributors | Involve them in reviews. Mentor them. Encourage them to become maintainers. |
| Build community around lessons | Quarterly online collaboration fest (CoFests) and Community calls. Chat (Matrix channel) |
| Publish periodically and recognize contributions | Author listed on tutorials. Hall of fame listing all contributors. Full tutorial citation at the end of the tutorial. Tweet about new or updated tutorials. List of new or updated tutorials in Galaxy Community newsletter. Soon: publication of tutorials via article |
| Evaluate lessons at several scales | Tutorial change (Pull Request) review. Embedded feedback form in tutorials for trainee feedback. Instructor feedback. Automatic workflow testing |
| Reduce, reuse, recycle | Sharing content between tutorials, specially using snippets. Development of small modular tutorials linked by learning paths |
| Link to other resources | Links to original paper and data, documentation, external tutorials and other material |
| You can't please everyone | but we can try (several different Galaxy introduction tutorials for different audience). Aim to clearly state what the tutorial does and does not cover, at the start. |

**S2 Table.** 10 Simple FAIR Rules. Implementation of the “10 simple rules for making training materials FAIR” [8] in the GTN training material framework

| Rules | Implementation in GTN framework |
| --- | --- |
| Plan to share your training materials online | Online training material portfolio (https://training.galaxyproject.org/), managed via a public GitHub repository (https://github.com/galaxyproject/training-material) |
| Improve findability of your training materials by properly describing them | Rich metadata associated with each tutorial that are visible and accessible via schema.org on each tutorial webpage |
| Give your training materials a unique identity | URL persistency with redirection in case of renaming of tutorials. Data used for tutorials stored on Zenodo and associated with a Digital Object Identifiers (DOI) |
| Register your training materials online | Tutorials automatically registered on TeSS, the ELIXIR’s Training e-Support System [41] |
| If appropriate, define access rules for your training materials | Online and free to use without registration |
| Use an interoperable format for your training materials | Content of the tutorials and slides written in Markdown. Metadata associated with tutorials stored in YAML, and workflows in JSON |
| Make your training materials (re-)usable for trainers | Online. Rich metadata associated with each tutorial: title, contributor details, license, description, learning outcomes, audience, requirements, tags/keywords, duration, date of last revision. Strong technical support for each tutorial: workflow, data on Zenodo and also available as data libraries on UseGalaxy.*, tools installable via the Galaxy Tool Shed, list of possible Galaxy instances with the needed tools. (Figure 3) |
| Make your training materials (re-)usable for trainees | Online and easy to follow tutorials. Rich metadata with “Specific, Measurable, Attainable, Realistic and Time bound” (SMART) learning outcomes following Bloom’s taxonomy. Requirements and follow-up tutorials to build learning path. List of Galaxy instances offering needed tools, data on Zenodo and also available as data libraries on UseGalaxy.*. Support chat embedded in tutorial pages. (Figure 3) |
| Make your training materials contribution friendly and citable | Open and collaborative infrastructure with contribution guidelines, a CONTRIBUTING file and a chat. Topic maintainers. How to cite tutorials and give credit to contributors available at the end of each tutorial (Figure 3). |
| Keep your training materials up-to-date | Open, collaborative and transparent peer-review and curation process. Short time between updates |

